## Supplemental Figures S1-4 for "Trade-offs between sperm viability and immune protein expression in honey bee queens (*Apis mellifera*)"

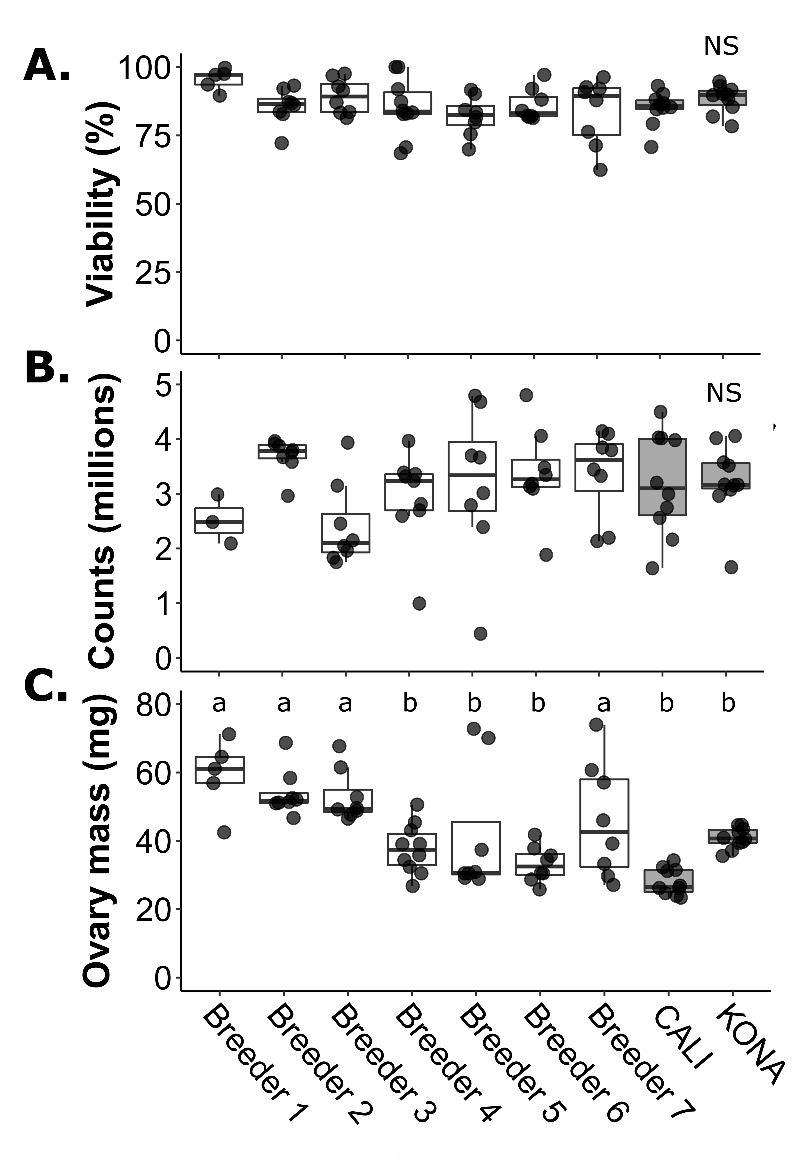

**Figure S1.** Data presented in **Figure 1**, separated by queen source. A) Sperm viability, B) sperm counts, and C) ovary mass metrics for healthy n = 52 healthy queens donated by 7 queen producers within British Columbia, n = 10 queens imported from California, and n = 10 queen imported from Hawaii. No differences in sperm viability nor counts were detected. Ovary masses varied significantly (one-way ANOVA), but this was likely an artifact of differences of caging time between producers. Lowercase letters indicate significant differences (p < 0.05).

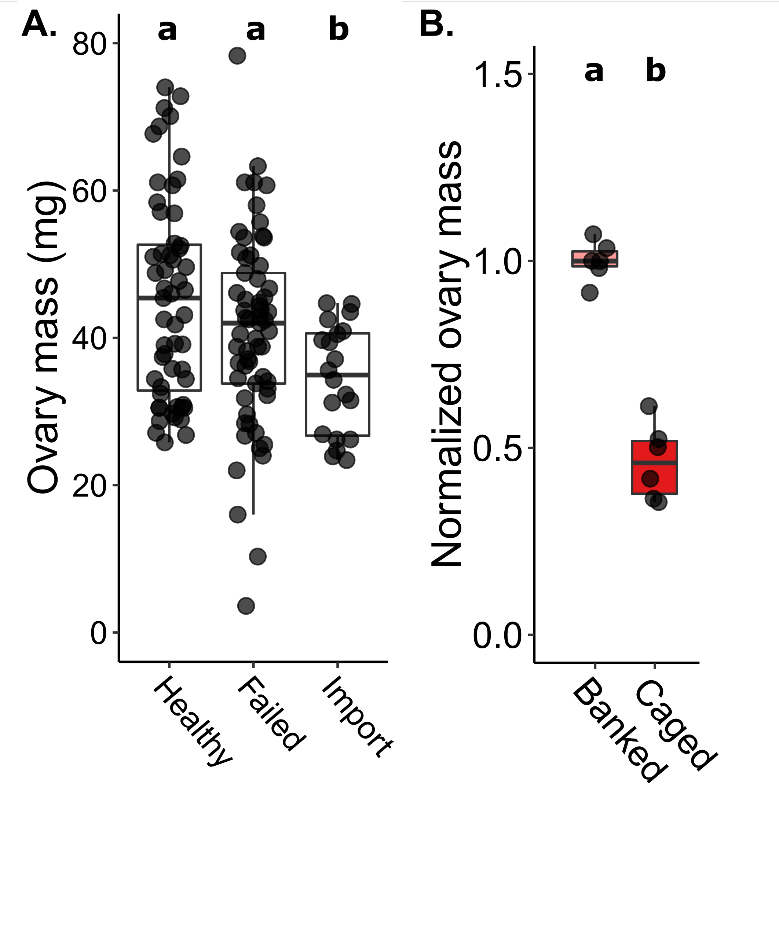

**Figure S2.** Queen ovary masses. A) Ovary mass data from healthy (n = 52), failed (n = 53), and imported (n = 20) queens (data also presented in **Figure 1**). Ovary mass of imported queens was significantly lower than healthy and failed. Lowercase letters indicate significant differences (p < 0.05). See **Table 1** for summary statistics. B) Ovary masses from imported Californian queens were either measured upon arrival (‘Caged,’ n = 6) or banked in a queenless colony (‘Banked,’ n = 6) for two weeks prior to measuring ovary mass. Lowercase letters indicate significant differences (t test, p < 0.05).

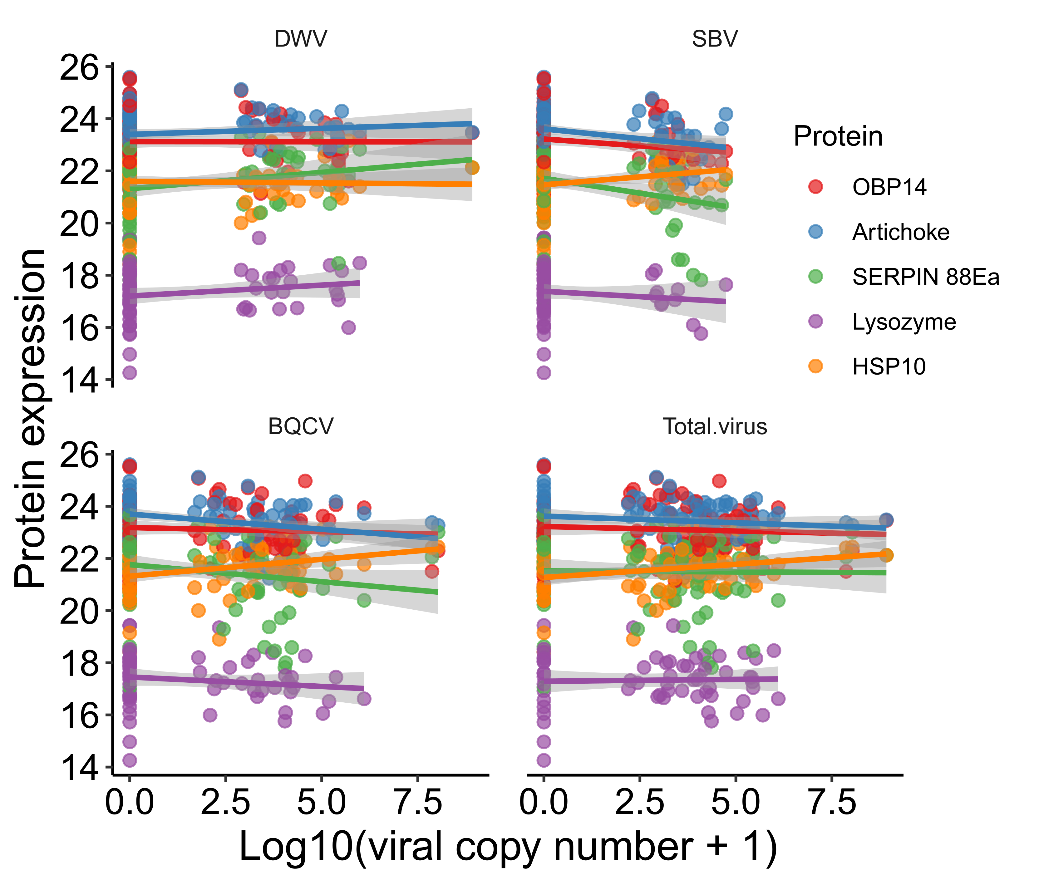

**Figure S3.** No associations between expression of top five proteins and viral factors. N = 94 queens had complete sets of viability, proteomics, and viral data. We evaluated relationships for each protein separately using a least squares linear model, including sperm viability, protein expression, queen status (levels: healthy, failed, imported), and viral copy numbers (levels: DWV, SBV, BQCV, and Total load) as fixed effects. No significant associations were identified (p > 0.05), except for protein expression with sperm viability, which we already determined previously (p < 0.005).

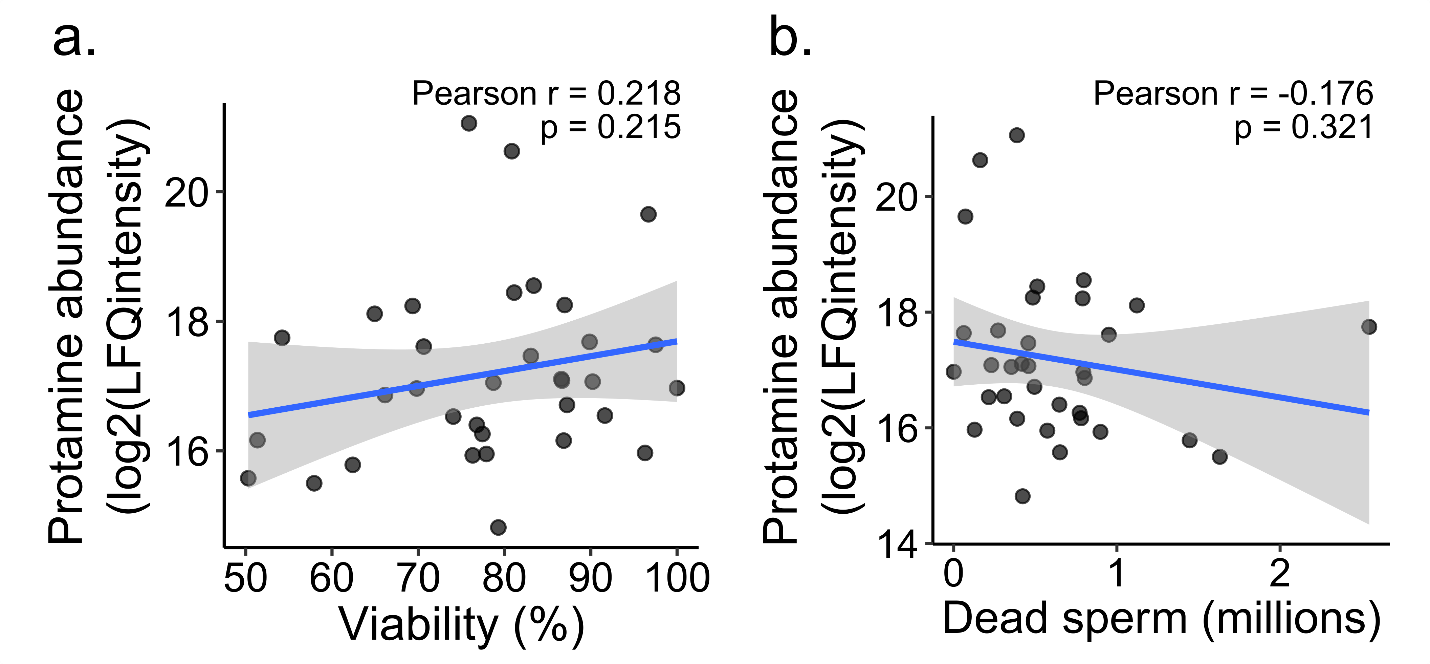

**Figure S4.** Correlations of Protamine-like protein (XP_026294833.1) with sperm viability (a) and number of dead sperm (b).
